# Endothelial struts, a mechanism to generate large lumenized blood vessels *de novo*

**DOI:** 10.1101/685180

**Authors:** Bart Weijts, Iftach Shaked, Wenqing Li, Mark Ginsberg, David Kleinfeld, David Traver

## Abstract

Lumenization of *de novo* formed blood vessels occurs either through cell hollowing (intracellular lumen)^1–3^ or cord hollowing (extracellular lumen)^4–6^ and restricts thereby the initial lumen diameter to one or two endothelial cells (ECs) respectively. However, vasculogenesis can result in large diameter blood vessels, raising the question how these vessels are formed. Here, we describe an alternative model of vasculogenesis that results in the formation of large diameter vessels. In this model, ECs coalesce into a branched network of EC struts within the future lumen of the vessel. These struts maintain the patency of the vessel and serve as a scaffold for the ECs forming the vessel wall, which initially consists out of a few patches of ECs. Together, we show that endothelial struts facilitate the formation of large blood vessels without being bound by the prerequisite of a cord-like structure, nor are they restricted in size.

---

*De novo* blood vessel formation occurs through the coalescence of ECs into a cord-like structure, subsequently followed by a cord-hollowing step. From *in vitro* and *in vivo* observations it has been shown that cord-hollow can occur through the fusion of intracellular lumens, formed through pinocytosis, of ECs connected in a head-to-tail orientation^1–3^. Recent studies, however, suggested that lumen formation more often occurs through the formation of an extracellular lumen between two ECs. This latter model depends on the apical-basal polarization of ECs, which leads to the rearrangements of cell-cell junctions to the periphery of the two adjacent ECs, delivery of negatively charged glycoproteins to the apical side of the cells and formation of a lumen between the two ECs through electrostatic repulsion^4,5^. Alternatively, this extracellular lumen might also be a result of apical exocytosis of vacuoles^6^. Depending on whether the lumen is formed intra- or extracellular, the initial diameter is restricted to 1 or 2 ECs respectively.

The dorsal aorta (DA) and posterior cardinal vein (PCV) are the first blood vessels to arise in the developing vertebrate embryo. The formation of the DA and PCV within the zebrafish model has been exemplary for studying the *de novo* formation of blood vessels^7,8^. In zebrafish, the caudal part of the PCV, posterior of the yolk sac extension, is referred to as the caudal vein (CV) and has the largest diameter of any blood vessel found during development. In perspective, the lumen diameter of the CV is about 5 times larger than that of the DA and forms through an extracellular lumen between two adjacent ECs^9–12^. Despite the large difference in lumen diameter between the DA and PCV, these vessels are formed and become functional in a near equal time-frame, suggesting that lumen of the CV might not be formed by one of the currently known mechanisms. To investigate how the CV is formed, we imaged its formation by confocal microscopy with high temporal and spatial resolution. For visualization purposes, we have distinguished between the formation of the lumen and the vessel wall by masking either population from the acquired images (Fig. 1a). Around 18 hours post fertilization (hpf) the venous ECs can be observed as single rounded cells at the midline of the embryo. Between 18-22 hpf, these ECs started to coalescence into a branching network of ECs that spans the future lumen of the CV (Fig. 1b; orange box and Supplementary Video S1 left panel). The majority of these branches are composed out of multiple ECs (Supplementary Fig. S1a). Within the next 4 hours this network gradually prunes and completely clears the lumen around 26-28 hpf (Fig. 1b-d and Supplementary Video S1 left panel). The wall of the CV exists initially only out of a few patches of ECs, which is in stark contrast with our current models of vasculogenesis in which the wall is already present before lumenization (Fig. 1b, green box and Supplementary Video S1 right panel). Closure of the vessel wall coincides with pruning of the endothelial network. Together, these results suggest that CV is formed around a branched network of ECs that serve both as a scaffold for the ECs forming the wall and as a temporary structure that maintains the patency of the lumen, hence we termed them endothelial struts. Bone morphogenetic protein (BMP) plays a pivotal role in the formation and remodeling of the caudal vein^13,14^. To test whether BMP signaling is involved in the formation of endothelial struts, we inhibited BMP signaling by either expressing the BMP antagonist *noggin*^14^ or by treatment with the small molecule DMH1, a dorsomorphin analogue^15^. *Noggin* expression was controlled by the heat-shock inducible promoter and allowed for inhibition of BMP signaling after the BMP-dependent specification of the dorsoventral mesoderm^16,17^. When we inhibited BMP signaling, the ECs maintained their rounded morphology and failed to coalescence into struts, indicating that BMP signaling is essential for endothelial strut formation (Fig. 1e). Pruning of the endothelial struts coincided with the closure of the wall of the CV (Fig. 1b). From these observations we hypothesized that ECs from endothelial struts are being re-used to form the wall of the CV. To test this hypothesis, we labeled and traced individual endothelial struts by ultraviolet photo-conversion of the green to red DENDRA2 fluorescent protein. We found that ECs from a single strut could be part of the wall of the CV, DA or give rise to the intersegmental vasculature. Thus, ECs from struts are being redistributed during pruning to become part of the surrounding vasculature (Supplementary Fig. S1b).

**Figure 1.**
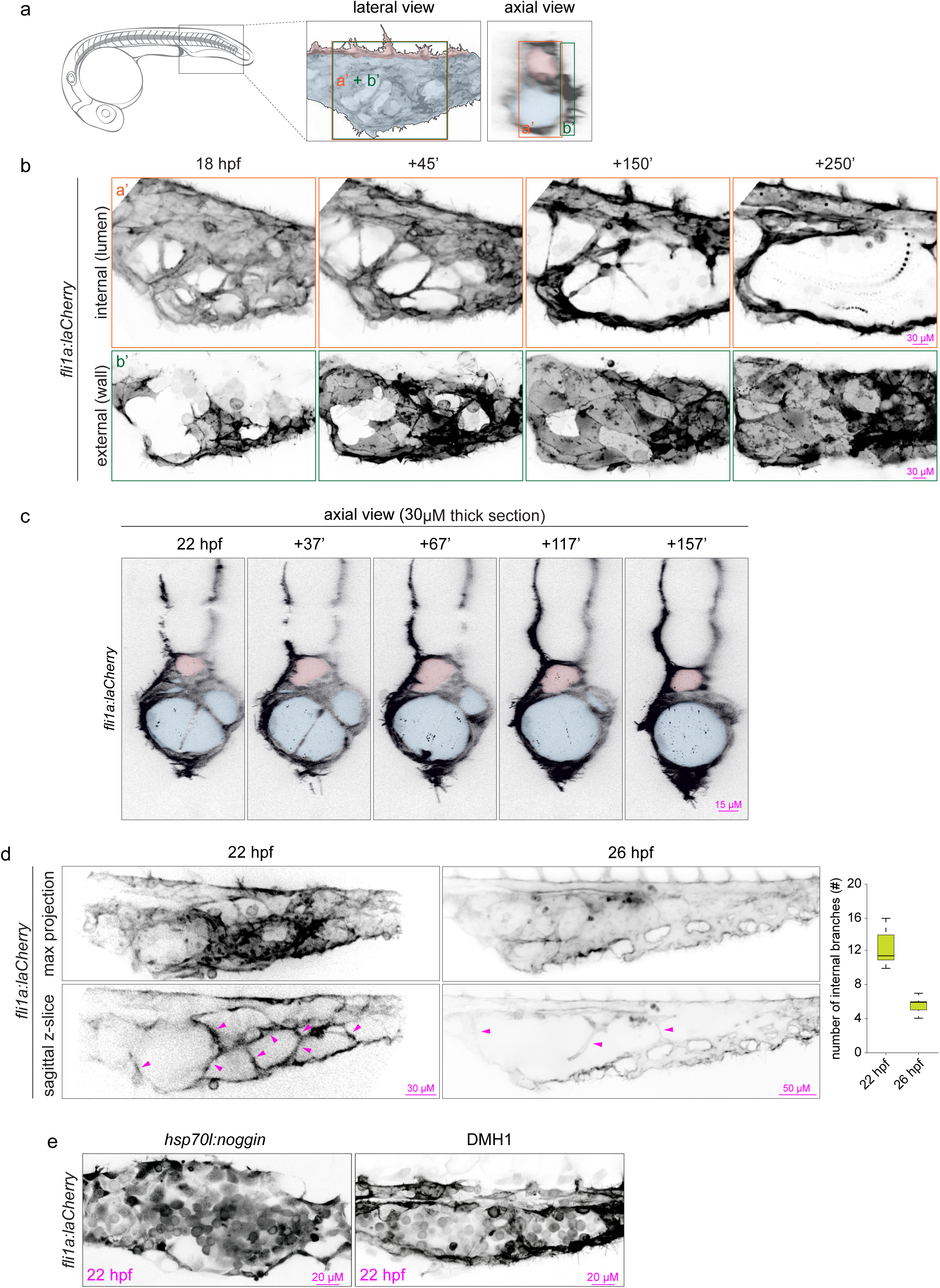
Formation of the caudal vein is facilitated through endothelial struts. (a) Schematic representation of a 24 hpf embryo; orange (lumen) and green (vessel wall) boxes indicate imaged regions. Lumen of the DA and CV are colored by red and blue respectively. (b) Orange panel; Stills from Supplementary Video S1 showing the formation and pruning of endothelial struts within the future lumen of the CV. Green panel; Stills from Supplementary Video S2 showing the formation of the vessel wall of the CV. (c) Stills of Supplementary Video S3 showing the axial view during the formation of the CV. Lumen of the DA and CV are colored by red and blue respectively. (d) Quantification of the number of endothelial struts at 22 hpf and 26 hpf by confocal microscopy and represented in a boxplot. Three independent experiments with a minimum of n = 25 animals per conditions per experiment were quantified. Arrow heads depict endothelial struts. (e) Inhibition of BMP signaling by heat-shock inducible expression of *noggin* (heats-hock at 14 hpf) or administration of 50 µM DMH1 from 18 hpf onwards.

It has been shown that the precursors of the arterial and venous ECs (also referred to as angioblasts) arise from distinct locations within the lateral plate mesoderm^11,12^. Due to this spatial difference, the arterial ECs arrive about 3 hours earlier at the midline of the embryo than the venous ECs and it was therefore hypothesized that the DA and PCV are formed independently by either arterial or venous ECs respectively^12^. Our finding that some endothelial struts located within the CV can be part of arterial blood vessels either suggests that venous ECs can give rise to arterial structures or arterial ECs participate in endothelial struts. To gain insight into the behavior of the arterial and venous EC populations, we first imaged the ECs populations when they arrived at the midline of the embryo and found that more anteriorly in the trunk the arterial and venous ECs can be observed as two separate populations (Fig. 2a). However, more posteriorly the arterial and venous ECs become increasingly intermingled without a clear separation between the two populations (Fig. 2a and 2b; left panel (18 hpf)). As a result, the DA and the CV form a common precursor vessel that overtime and at the interface of the DA and CV, separates into two separate vessels through unmixing of arterial and venous ECs (Fig. 2b and Supplementary Video S2). To visualize arterial and venous ECs, we imaged embryos in which arterial ECs are fluorescently labeled by the arterial restricted Notch ligand *delta-like 4* (*dll4*)^18^ or by fluorescent reporting of Notch activity (TP1)^19^. Some endothelial struts located with the CV express arterial markers (Fig. 2c and d). Notably, we observed that arterial ECs that form the roof of the DA -juxtaposed the hypochord-expressed *dll4* earlier and showed an increased Notch signaling compared to the arterial ECs positioned more ventrally (Fig. 2c and d). These results suggest that the arterial angioblast positioned against the hypochord have a stronger arterial signature than the more ventrally positioned arterial ECs, which includes the ones that form endothelial struts. The EphrinB2a transmembrane ligand is a downstream target of Notch signaling and plays a key role in determining arterial and venous boundaries and segregation^10,20,21^. Inhibiting EphrinB2a signaling prevented the unmixing of the arterial and venous ECs in the CV region, resulting in a DA and PCV that remained fused (Fig. 2e)^10^. However, abrogation of *ephrinB2a* expression did not prevent pruning of endothelial struts (Fig. 2e). Thus, the most ventral arterial ECs have an initial weaker arterial identity that becomes stronger over time, driving *ephrinB2a* expression and thereby the segregation of the two vessels, while it is not involved in pruning of the endothelial struts.

**Figure 2.**
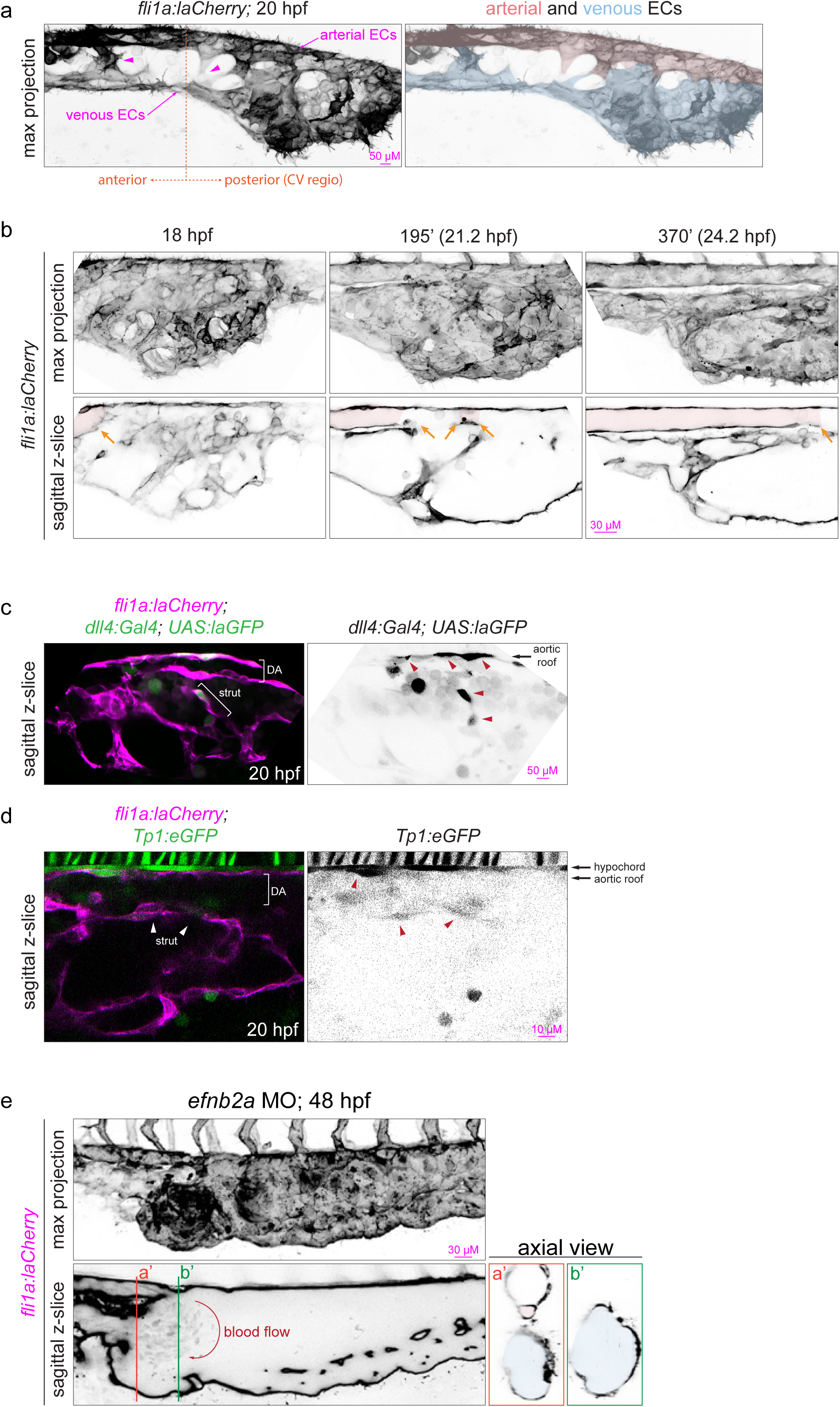
The CV arises from a common pre-cursor vessel. (a) Arterial and venous EC populations. (b) Stills from Video S3, the CV and DA are formed through segregation of arterial and venous ECs from a common pre-cursor vessel. Red color and arrows depict the expanding lumen of the DA. (c) Expression of the arterial restricted notch ligand allows for the discrimination of arterial (green) and venous ECs (magenta, all ECs). (d) Similar to C except that the arterial ECs are marked by GFP which is under the control of a Notch responsive promoter (TP1; green). (E) Morpholino mediated knock down of *ephrinB2* prevents the segregation of arterial and venous ECs.

Pruning of endothelial struts coincides with the onset of blood flow, an important factor in vascular remodeling and endothelial cell fate determination^22–24^. Therefore, we tested whether pruning of endothelial struts was dependent on blood flow. To this end, we prevented the onset of blood flow by inhibiting the expression of *cardiac troponin T2a* (*tnnt2a*) and thereby the myocardial function or we administrated the muscle relaxant ms-222 (tricaine methanesulfonate) to suppress the heart beat^25^. To monitor blood flow, we treated embryos in which primitive erythrocytes and the ECs are differently labeled and found that embryos without blood flow showed no difference in endothelial strut pruning (Fig. 3a). Our imaging data revealed that the primitive erythrocytes were positioned between the arterial and venous ECs, with an increased mixing between the erythrocytes and venous ECs from anterior to posterior (Fig. 3c). As struts are formed, the CV becomes compartmentalized with some compartments filled with erythrocytes (Fig. 3c). The DA is initially devoid of erythrocytes (Fig. 3b and c (20 hpf)) and only start to enter the DA when blood flow is initiated (Fig. 3c (24 hpf)). Pruning of the endothelial struts resulted in the gradual loss of these compartments, thereby releasing the erythrocytes into the circulation (Fig. 3d and Supplementary Video S3).

**Figure 3.**
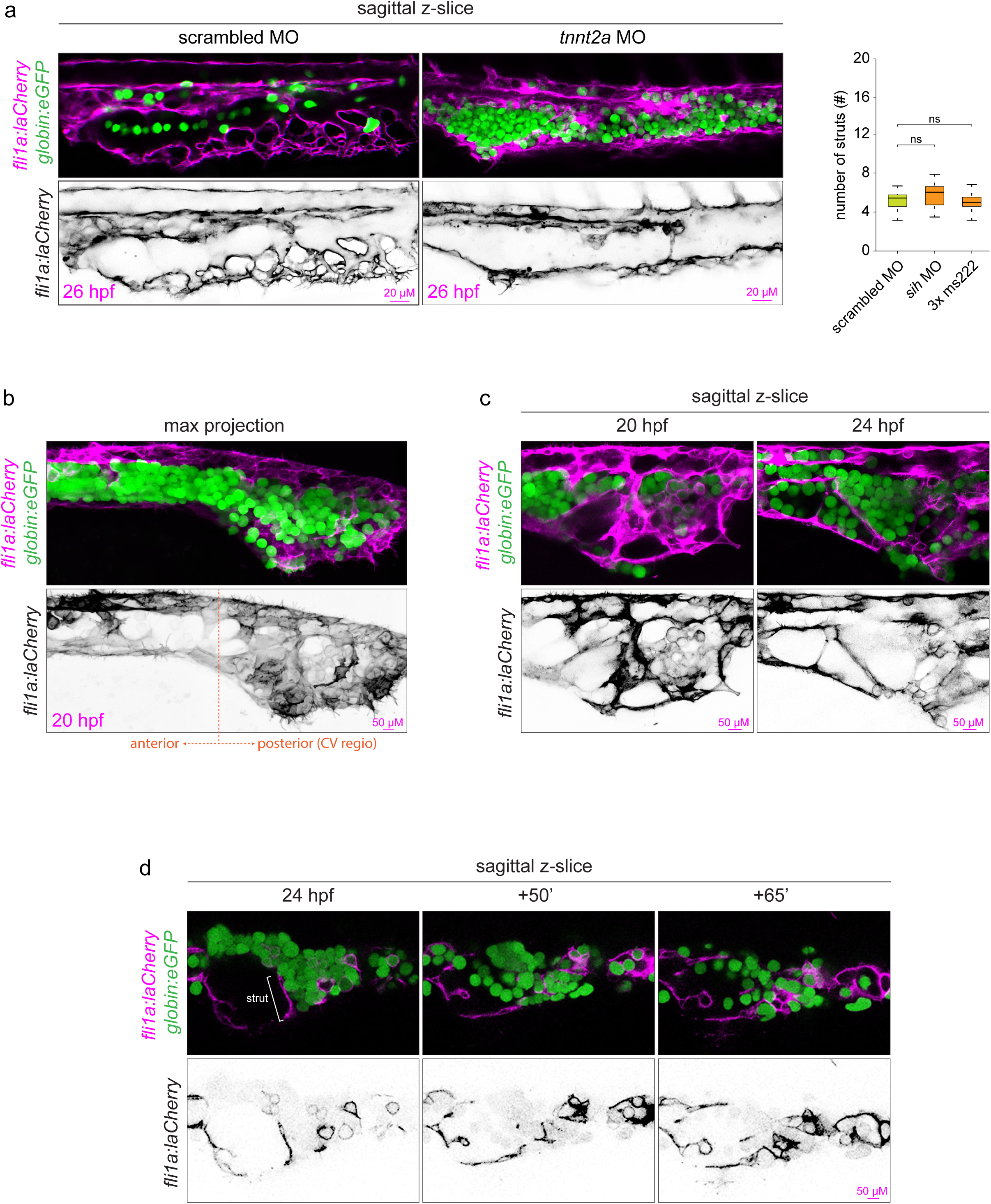
Endothelial struts compartmentalize the CV. (a) Onset of blood flow was prevented by morpholino knock down of *tnnt2a* or by administration of 3 times concentrated ms222 (∼0.06%). Primitive erythrocytes are marked in green. Endothelial struts were quantified and data represented in a boxplot. Three independent experiments with a minimum of n = 25 animals per conditions per experiment were quantified. ns = not significant. (b) Primitive erythrocytes (green) are positioned between the arterial and venous ECs, with an increased mixing of the population from anterior to posterior. (c) Endothelial struts compartmentalize the CV, with some compartments filled with erythrocytes (green). (d) Stills of Supplementary Video S4, pruning of the indicated strut results in the release of the erythrocytes into the circulation.

To study the function of endothelial struts, we ablated 1-2 cells within a single strut by ultrashort pulses of near-infrared laser light that severed the strut. This technique has been show to generate negligible heat transfer and collateral damage to neighboring tissues^26^. Due to the anatomical structure of the CV, we could only reliably sever endothelial struts after 22-24 hpf (Fig. 4 and Supplementary Video S4). Struts perpendicular to the sagittal plane were often difficult to detect and thus reliably to sever, but on average we were able to ablate 95% of the struts (Fig. 4; arrow head middle right panel and Supplementary Video S5). As a control, we sham treated embryos by aiming the laser 25-50 micron next to endothelial struts, in the empty space of the lumen, with the same number of ablation pulses as in treated embryos. Severing of nearly all endothelial struts did initially not lead to a dramatic change in the shape of the lumen. However, the CV collapsed upon the onset of circulation, which can be explained by the presence of the primitive erythrocytes, which hold the shape of the CV until they are flushed from the CV by circulation. (Fig. 4b and Supplementary Video S6). Nonetheless, the lumen of the collapsed CV remained sufficiently large to allow circulation. Severing a single strut often resulted in a slight deformation of the CV and in animals in which circulation just had initiated it also resulted in the release of erythrocytes into the circulation. (Supplementary Fig. S2 and Supplementary Video S5 (embryo 2)). Normally, the CV remodels in to the caudal vein plexus through sprouting and anastomosis, a blood flow dependent process^27–29^. However, embryos devoid of struts showed an impaired remodeling of the CV, which might be due to altered flow strength and patterns (Fig. 4c).

**Figure 4.**
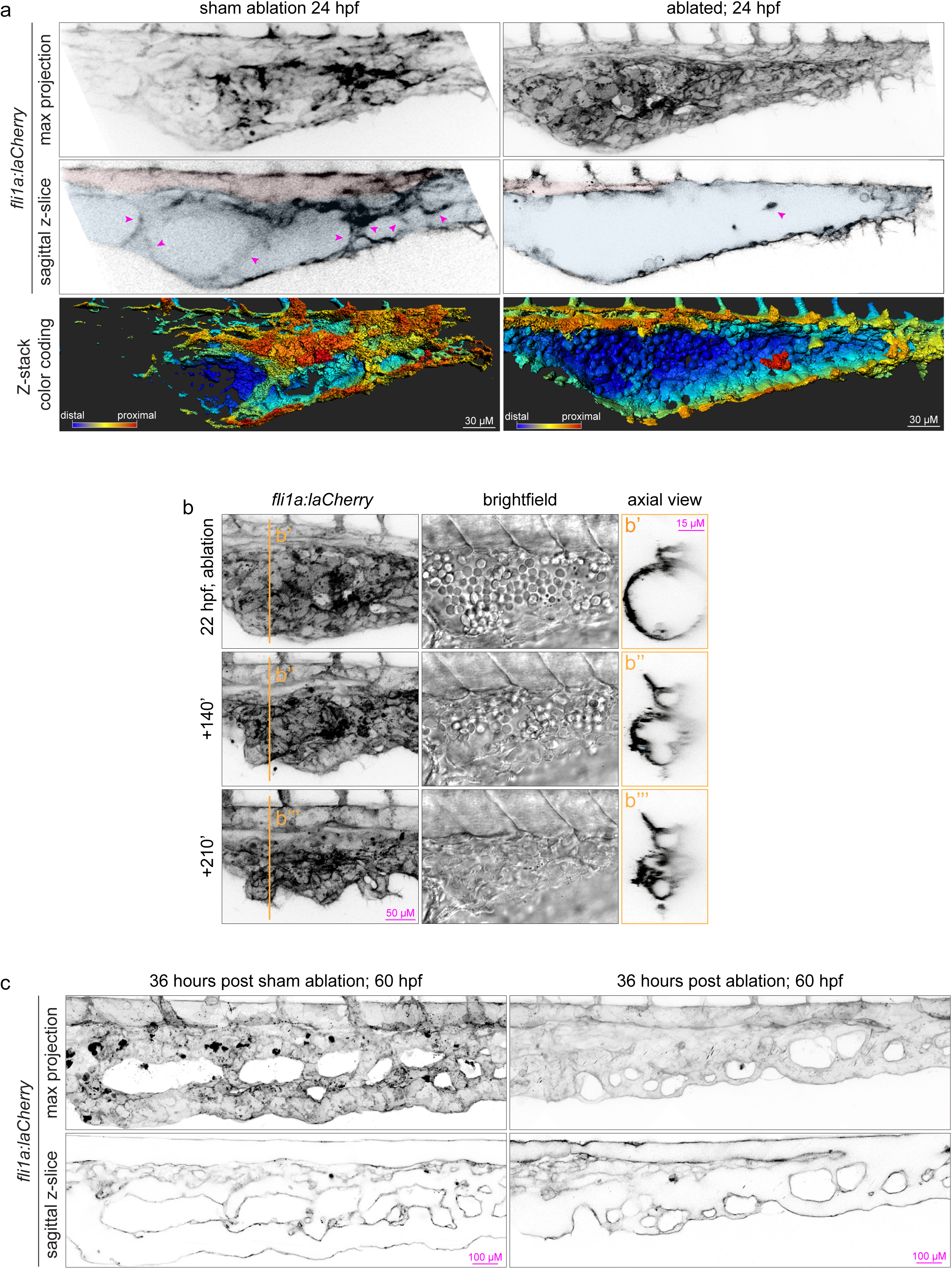
Endothelial struts provide structural support to the CV. (a) Laser ablation of endothelial struts. In sham treated animals, the laser was fired ∼25-50 micron next to the strut in the empty space of the lumen, with the same number of ablation pulses as in treated embryos. Ablation was carried out with 25 pulses of 0.2μJ delivered at 5kHz (b) Stills of Supplementary Video S6, ablation of endothelial struts results in the collapse of the CV after initiation of circulation. (c) The CV without endothelial struts shows defects in remodeling 36 hours post ablation.

Here, we show that during embryonic development ECs can form large diameter blood vessels almost instantaneously through the formation of endothelial struts, without the prerequisite of a vascular cord (Fig. 5). Severing endothelial struts results in the collapse of the CV upon the onset of circulation suggesting that these struts provide structural support and maintain the shape of the lumen by withstanding external forces. In support, severing a single endothelial strut resulted in a slight compression of the CV (Supplementary Fig. S2), suggesting that endothelial struts experience a compressional force. These forces are generated during development and are an import factor in tissue morphogenesis and patterning^30,31^. However, most endothelial struts are as thin as 1 or 2 ECs and therefore it is arguable whether they can withstand large external forces and provide structural support directly. Combinatory imaging of the primitive erythrocytes and endothelial struts revealed that endothelial struts compartmentalize the CV, of which some are filled with erythrocytes. We propose that these erythrocyte filled compartments provide additional structural support and completement thereby the endothelial struts in maintaining the shape of the CV.

**Figure 5.**
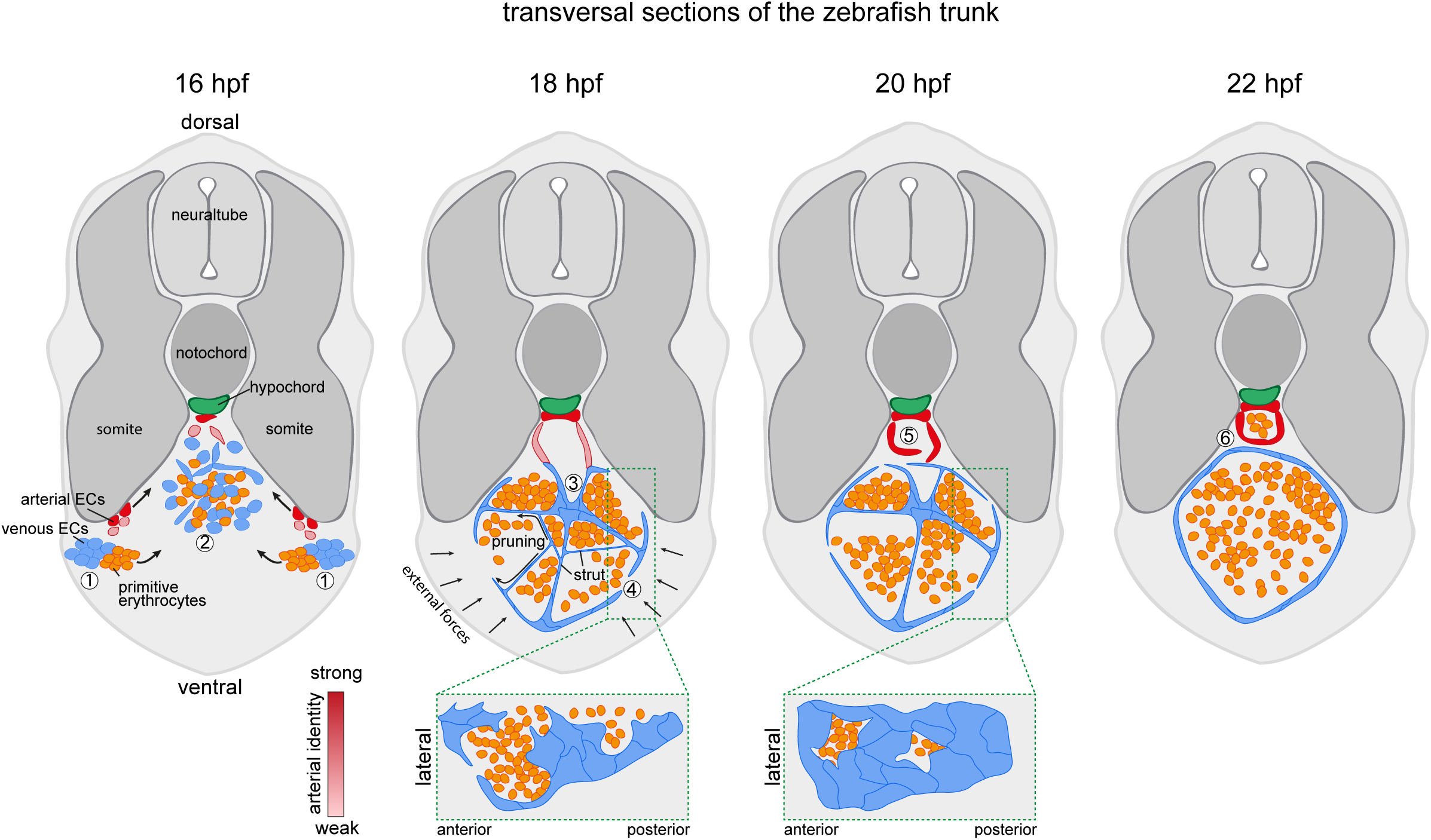
Endothelial strut model. 1) Venous (blue) and arterial (red) EC precursors originate from distinct locations within the lateral plate mesoderm, with the primitive erythrocytes (precursors) positioned medially adjacent to the venous ECs. 2) EC precursors and primitive erythrocytes migrate towards the midline of the embryo. Arterial ECs migrate along the ventral face of the somites and arterial ECs in direct contact with the somites receive inductive cues that strengthen the arterial fate (shades of red). 3) At the midline, venous ECs start to branch and form a dense network of endothelial struts, these struts compartmentalize the caudal vein. Since venous ECs are mixed with the primitive erythrocytes, the formation of struts encloses the primitive erythrocytes in compartments. Furthermore, at this point the DA and PCV exist as one large precursor vessel with some struts formed by both arterial and venous ECs, these arterial ECs have a weak arterial identity. 4) The vessel wall consists initially only out of a few patches of ECs. Upon pruning of the endothelial struts, the ECs from the strut migrate into the vessel wall, which gradually closes. 5) Segregation of the precursor vessel into the DA and PCV occurs through unmixing of arterial and venous ECs, a process that is dependent on Ephrin signaling. Laser ablation of endothelial struts results into the collapse of the lumen suggesting that struts maintain the patency of the vessel by withstanding external forces. 6) Struts have completely cleared the lumen of the CV which is maintained by blood pressure.

It was recently shown that the lateral plate mesoderm harbors two populations of angioblasts, the most medial population -closest to the midline of the embryo-forms the DA and the more lateral population forms the PCV^11,12^. Due to this spatial difference, the arterial angioblasts arrive about 2-3 hours earlier at the midline and are positioned ventrally against the hypochord, while the venous angioblast migrate more ventrally of the arterial population. From these observations it was concluded that the DA and PCV arise as two separate entities ^12^ and not through a previous proposed model of arterial-venous segregation^10^. In this latter model, angioblasts first form a common pre-cursor vessel at the midline of the embryo that subsequently segregates into the DA and PCV through unmixing of arterial and venous ECs, a process dependent on EphrinB2a^10^. While these two models seem mutual exclusive, we show here that the caudal part of the DA and CV first forms a common pre-cursor vessel through endothelial struts and subsequently segregates into the DA and PCV in an EphrinB2a dependent mechanism. However, more anteriorly the ECs appear as two separate populations that sporadically interact with each other, suggesting that the DA and PCV arise as two separate vessels (Fig. 2a) To date, it remains unclear how and precisely when the angioblasts are specified into arterial or venous ECs. However, it has become clear that angioblasts committing to the arterial fate require multiple spatial and temporal Notch signal inputs, suggesting that angioblast gradually acquire the arterial identity^12,32,33^. This gradual specification plays a key role in endothelial strut formation as it allows firstly for the development of endothelial struts between ECs of the opposite fate and secondly it facilitates the unmixing of arterial and venous ECs through expression of arterial and venous specific genes like EphrinB2 when the arterial identity becomes more prominent^10,21^. Together, we show here that depending on the anatomical location in the trunk the DA and PCV can either arise as separate vessels or from a common pre-cursor vessel through endothelial strut formation.

Many organisms rely on the formation of tubes to transport fluids or gasses, hence the great variation in shapes, sizes and composition. Large tubes are generally thought to arise through wrapping or cavitation, which requires massive cell reorganization or loss through apoptosis respectively^34^. The here presented mechanism provides a new concept to form such large lumenized structures without the need of a prerequisite sheet of cells (wrapping) or the unnecessary loss of a large number of ECs.

## Methods

### Zebrafish husbandry

Zebrafish (Danio rerio) were maintained according to the guidelines of the UCSD Institutional Animal Care and Use Committee. The following zebrafish lines have been previously described: *Tg(fli1a:lifeactCherry)*^*ncv7Tg* 27^ referred to as fli1a:laCherry; *Tg(fli1a:eGFP)*^*y1* 35^; *Tg(hbbe1*.*1:EGFP)*^*zf446* 36^ referred to as *Tg(globin:eGFP); Tg(flk:DENDRA2)*^37^; *Tg(dll4:Gal4FF)*^*hu10049Tg* 18^; *Tg(UAS:lifeactGFP)*^*mu271* 38^ referred to as UAS:laGFP; *Tg(EPV*.*Tp1-Mmu*.*Hbb:GFP-utrn)* ^19^ referred to as *Tg(Tp1:GFP*).

### Morpholino, plasmid injections, heat-shock and chemical treatment

Embryos were injected at the one-cell stage with 2.5 nl morpholino oligonucleotides (MOs) or 50 ng plasmid with 100ng tol2 mRNA. EphrinB2a (efnb2a) translation blocking MO (GeneTools) (5’-CGGTCAAATTCCGTTTCGCGGGA −’) at 2.5 ng/nl^39^; Silent heart morpholino *tnnt2a* (5’-CATGTTTGCTCTGATCTGACACGCA −3’) at 2 ng/nl ^25^. Capped tol2 mRNA was synthesized from linearized pCS2+ constructs using the mMessage mMachine SP6 kit (Ambion, AM1340). Injected and un-injected control embryos were heat-shocked at 38 °C for 30 minutes. Embryos were treated with 50 µM DMH1 dissolved in DMSO (1000X stock solution) and control embryos were treated with DMSO alone.

### Microscopy and laser ablation

Live microscopy was done in environmentally controlled microscopy systems based on a Leica TSP LSM 5 confocal microscope, a Leica TCS SP8 DLS or a Zeiss LSM 880 Airyscan. For all imaging, embryos were placed into a modified Four-Well WillCo dish ^40^ submerged in E3 medium containing 0.168 mg/ml = 0.0168% = ∼0.02% Ethyl 3-aminobenzoate methanesulfonate (MS222; sigma E10521) at a temperature of 28.5 °C. Imaging was subsequently done with either 20×/0.75, 40×/1.4 or 63×/1.46 objectives.

For laser ablation experiments, embryos were embedded in 1% low melting point agarose (Invitrogen) in a WillCo dish with a 40 mm #1.5 cover glass bottom and placed under a two-photon laser scanning microscope of local design that included an amplified beam^41^. Laser ablation of endothelial struts was achieved by using targeted ultra-fast laser pulses that were generated with a multi-pass Ti:Al2O3 amplifier of local construction that followed a previously published design^26^ and operated at a 5 kHz pulse-rate. The Ablation beam and the imaging beam were combined before the microscope objective with a polarizing beam splitter^26^. We focused the two beams in the same focal plane and centered the ablation beam in the area that is raster-scanned by the imaging beam so that ablation occurred at the center of the TPLSM imaging field. The energy per pulse of the ablation beam was tuned with neutral density filters and the number of pulses was controlled by a mechanical shutter (Uniblitz LS3Z2 shutter and VMM-D1 driver; Vincent). The energy and number of pulses was modified based on damage assessed from the real-time TPLSM images. Ablation was carried out with 25 pulses of 0.2µJ delivered at 5 kHz

### Statistical analysis

For each in vivo experiment, animals from the same clutch were divided into different treatment groups without any bias. The whole clutch was excluded if more than 10% of control embryos displayed obvious developmental defects. At least 20 animals from each treatment group were randomly picked for analysis. At least three independent experiments were performed per each treatment group. For in vitro experiments, at least three independent experiments were performed per each condition. Statistical analysis was performed using SPSS 20 (IBM). Mann–Whitney U test was used for statistical analysis of two groups, unequal variances. Unpaired t-test was used for two groups, equal variances. Kruskal-Wallis test was used for statistical analysis of multiple groups, equal variances, and 1-way ANOVA, for multiple groups, unequal variances. Dunns post-hoc test was used for pairwise multiple comparisons. P values < 0.05 were considered significant.

## Acknowledgments

We thank Professor Debby Yelon (University of California San Diego) and Suk-Won Jin (Yale Cardiovascular Research Center) for generously providing us with the silent heart morpholino and hsp70l:noggin plasmid, respectively. Dr. Phil Tsai for use of the Q-bio laboratory confocal. We thank Jennifer Santini, Neeraj Gohad (Zeiss) and Kristofer Fertig (Leica) for microscopy technical assistance and the UCSD School of Medicine Microscopy Core. This work was supported by the San Diego School of Medicine Microscopy Core (P30 NS047101)

## Author contributions

B.W. and D.T. designed and analyzed experiments; B.W. performed all experiments; Laser ablation experiments were performed by B.W. and I.S under supervision of D.K. B.W. and D.T. wrote the manuscript.

## Competing interests

The authors declare no competing financial interests.

**Supplementary Figure S1.**
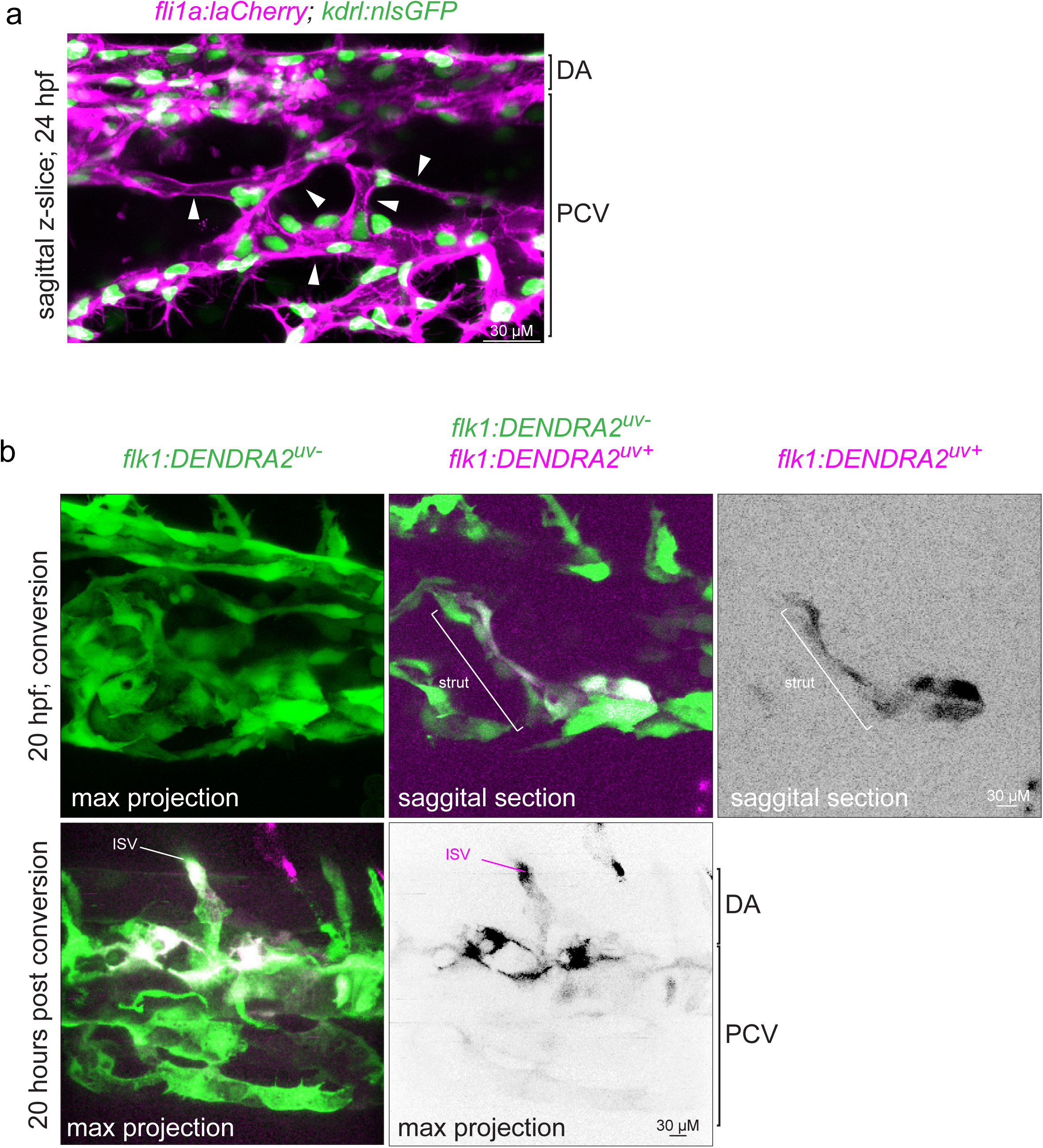
Characterization of endothelial struts. (a) ECs nuclei (green) show that endothelial branches within the CV consists out of multiple ECs. Arrow heads point to single endothelial branches. (b) Lineage tracing of single endothelial struts by photo-conversion of DENDRA2 (green-to-red).

**Supplementary Figure S2.**
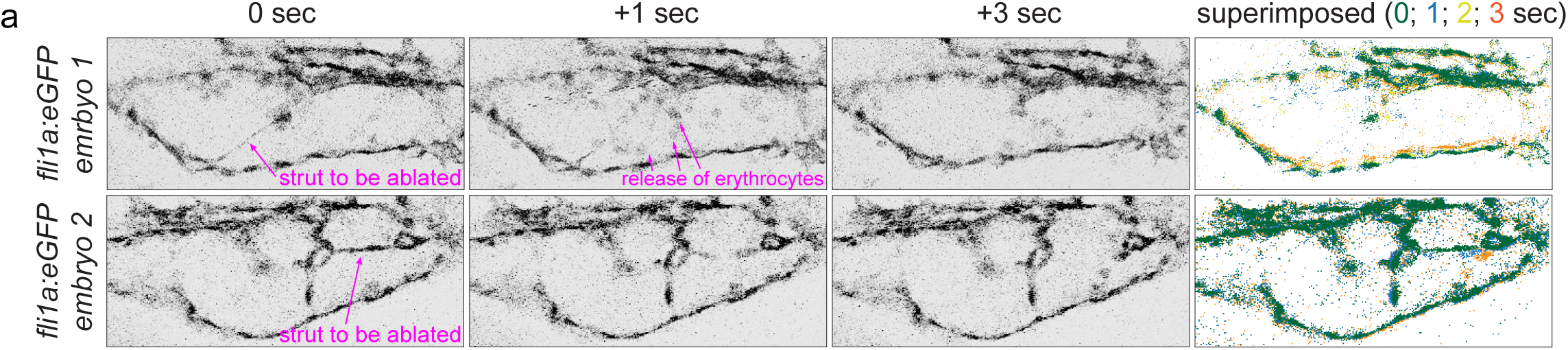
Endothelial strut ablation. Stills of Supplementary Video S6 and S7 show single strut ablation. Superimposed images show that ablation of a single strut can result in deformation of the CV shape.

Supplementary Video S1, formation of the caudal vein

Supplementary Video S2, the dorsal aorta and caudal vein are formed from a common pre-cursor vessel

Supplementary Video S3, pruning of endothelial struts results in the gradual release of primitive erythrocytes into circulation

Supplementary Video S4, laser ablation of endothelial struts Supplementary Video S5, visualization of the caudal vein post ablation

Supplementary Video S6, endothelial struts maintain the shape of the caudal vein lumen

## References

1. Folkman, J. & Haudenschild, C. Angiogenesis in vitro. Nature 288, 551–556 (1980).

2. Kamei, M. et al. Endothelial tubes assemble from intracellular vacuoles in vivo. Nature 442, 453–456 (2006).

3. Davis, G. E. & Bayless, K. J. An integrin and Rho GTPase-dependent pinocytic vacuole mechanism controls capillary lumen formation in collagen and fibrin matrices. Microcirculation 10, 27–44 (2003).

4. Strilić, B. et al. Electrostatic cell-surface repulsion initiates lumen formation in developing blood vessels. Curr Biol 20, 2003–2009 (2010).

5. Ferrari, A., Veligodskiy, A., Berge, U., Lucas, M. S. & Kroschewski, R. ROCK-mediated contractility, tight junctions and channels contribute to the conversion of a preapical patch into apical surface during isochoric lumen initiation. J Cell Sci 121, 3649–3663 (2008).

6. Blum, Y. et al. Complex cell rearrangements during intersegmental vessel sprouting and vessel fusion in the zebrafish embryo. Dev Biol 316, 312–322 (2008).

7. Hogan, B. M. & Schulte-Merker, S. How to plumb a pisces: understanding vascular development and disease using zebrafish embryos. Dev Cell 42, 567–583 (2017).

8. Gore, A. V., Monzo, K., Cha, Y. R., Pan, W. & Weinstein, B. M. Vascular development in the zebrafish. Cold Spring Harb Perspect Med 2, a006684 (2012).

9. Strilić, B. et al. The molecular basis of vascular lumen formation in the developing mouse aorta. Dev Cell 17, 505–515 (2009).

10. Herbert, S. P. et al. Arterial-venous segregation by selective cell sprouting: an alternative mode of blood vessel formation. Science 326, 294–298 (2009).

11. Jin, S.-W., Beis, D., Mitchell, T., Chen, J.-N. & Stainier, D. Y. R. Cellular and molecular analyses of vascular tube and lumen formation in zebrafish. Development 132, 5199–5209 (2005).

12. Kohli, V., Schumacher, J. A., Desai, S. P., Rehn, K. & Sumanas, S. Arterial and venous progenitors of the major axial vessels originate at distinct locations. Dev Cell 25, 196–206 (2013).

13. Heinke, J. et al. Antagonism and synergy between extracellular BMP modulators Tsg and BMPER balance blood vessel formation. J Cell Sci 126, 3082–3094 (2013).

14. Wiley, D. M. et al. Distinct signalling pathways regulate sprouting angiogenesis from the dorsal aorta and the axial vein. Nat Cell Biol 13, 686–692 (2011).

15. Hao, J. et al. In vivo structure-activity relationship study of dorsomorphin analogues identifies selective VEGF and BMP inhibitors. ACS Chem Biol 5, 245–253 (2010).

16. Mullins, M. C. et al. Genes establishing dorsoventral pattern formation in the zebrafish embryo: the ventral specifying genes. Development 123, 81–93 (1996).

17. Kishimoto, Y., Lee, K. H., Zon, L., Hammerschmidt, M. & Schulte-Merker, S. The molecular nature of zebrafish swirl: BMP2 function is essential during early dorsoventral patterning. Development 124, 4457–4466 (1997).

18. Hermkens, D. M. A. et al. Sox7 controls arterial specification in conjunction with hey2 and efnb2 function. Development 142, 1695–1704 (2015).

19. Parsons, M. J. et al. Notch-responsive cells initiate the secondary transition in larval zebrafish pancreas. Mech Dev 126, 898–912 (2009).

20. Adams, R. H. et al. Roles of ephrinB ligands and EphB receptors in cardiovascular development: demarcation of arterial/venous domains, vascular morphogenesis, and sprouting angiogenesis. Genes Dev 13, 295–306 (1999).

21. Wang, H. U., Chen, Z. F. & Anderson, D. J. Molecular distinction and angiogenic interaction between embryonic arteries and veins revealed by ephrin-B2 and its receptor Eph-B4. Cell 93, 741–753 (1998).

22. Chen, Q. et al. Haemodynamics-driven developmental pruning of brain vasculature in zebrafish. PLoS Biol 10, e1001374 (2012).

23. Weijts, B. et al. Blood flow-induced Notch activation and endothelial migration enable vascular remodeling in zebrafish embryos. Nat Commun 9, 5314 (2018).

24. Le Noble, F. et al. Flow regulates arterial-venous differentiation in the chick embryo yolk sac. Development 131, 361–375 (2004).

25. Sehnert, A. J. et al. Cardiac troponin T is essential in sarcomere assembly and cardiac contractility. Nat Genet 31, 106–110 (2002).

26. Nishimura, N. et al. Targeted insult to subsurface cortical blood vessels using ultrashort laser pulses: three models of stroke. Nat Methods 3, 99–108 (2006).

27. Wakayama, Y., Fukuhara, S., Ando, K., Matsuda, M. & Mochizuki, N. Cdc42 mediates Bmp-induced sprouting angiogenesis through Fmnl3-driven assembly of endothelial filopodia in zebrafish. Dev Cell 32, 109–122 (2015).

28. Phng, L.-K., Stanchi, F. & Gerhardt, H. Filopodia are dispensable for endothelial tip cell guidance. Development 140, 4031–4040 (2013).

29. Karthik, S. et al. Synergistic interaction of sprouting and intussusceptive angiogenesis during zebrafish caudal vein plexus development. Sci Rep 8, 9840 (2018).

30. Heisenberg, C.-P. & Bellañche, Y. Forces in tissue morphogenesis and patterning. Cell 153, 948–962 (2013).

31. Narciso, C. E., Contento, N. M., Storey, T. J., Hoelzle, D. J. & Zartman, J. J. Release of applied mechanical loading stimulates intercellular calcium waves in drosophila wing discs. Biophys J 113, 491–501 (2017).

32. Kobayashi, I. et al. Jam1a-Jam2a interactions regulate haematopoietic stem cell fate through Notch signalling. Nature 512, 319–323 (2014).

33. Quillien, A. et al. Distinct Notch signaling outputs pattern the developing arterial system. Development 141, 1544–1552 (2014).

34. Lubarsky, B. & Krasnow, M. A. Tube Morphogenesis. Cell 112, 19–28 (2003).

35. Lawson, N. D. & Weinstein, B. M. In vivo imaging of embryonic vascular development using transgenic zebrafish. Dev Biol 248, 307–318 (2002).

36. Ganis, J. J. et al. Zebrafish globin switching occurs in two developmental stages and is controlled by the LCR. Dev Biol 366, 185–194 (2012).

37. Tian, Y. et al. The first wave of T lymphopoiesis in zebrafish arises from aorta endothelium independent of hematopoietic stem cells. J Exp Med 214, 3347–3360 (2017).

38. Helker, C. S. M. et al. The zebrafish common cardinal veins develop by a novel mechanism: lumen ensheathment. Development 140, 2776–2786 (2013).

39. Cooke, J. E., Kemp, H. A. & Moens, C. B. EphA4 is required for cell adhesion and rhombomere-boundary formation in the zebrafish. Curr Biol 15, 536–542 (2005).

40. Weijts, B., Tkachenko, E., Traver, D. & Groisman, A. A Four-Well Dish for High-Resolution Longitudinal Imaging of the Tail and Posterior Trunk of Larval Zebrafish. Zebrafish 14, 489–491 (2017).

41. Tsai, P. S. & Kleinfeld, D. in In Vivo Optical Imaging of Brain Function (eds. Frostig, R. D., Frostig, R. D., Frostig, R. D. & Frostig, R. D.) (CRC Press/Taylor & Francis, 2009).

